# Modeling platinum resistance in a stem-like patient-derived ovarian cancer sample

**DOI:** 10.1101/2024.01.30.577975

**Authors:** Tise Suzuki, Ashlyn Conant, Yeonkyu Jung, Ryan Bax, Ashley Antonissen, Wanqiu Chen, Gary Yu, Yevgeniya J. Ioffe, Charles Wang, Juli J. Unternaehrer

## Abstract

**Background:** Chemoresistance and tumor recurrence remain a significant challenge in ovarian cancer. Particularly in the context of platinum resistance, many mechanisms have been identified, including the activation of cellular processes like epithelial-mesenchymal transition (EMT), which generates cells with stemness characteristics. Current models of platinum resistance are limited or not adequate representations of the heterogeneity of the disease. Thus, to advance our understanding of chemoresistance in the context of cancer stem cells (CSC) in ovarian cancer, this study aims to develop an effective model for cisplatin resistance using a patient-derived cancer stem-like sample.

**Methods:** PDX4, a patient-derived cancer cell line with stem-like properties, was exposed to increasing concentrations of cisplatin *in vitro* in parallel with vehicle treated cells. Once chemoresistance was established and confirmed, the resistance model was validated through comprehensive molecular profiling through RNA- and miRNA-sequencing, followed by the assessment of alterations in cell morphology, protein expression, and functional properties in the context of EMT and cancer stemness. Moreover, we explored potential signaling pathways involved in cisplatin resistance in these stem-like cancer cells.

**Results:** Our findings reveal the presence of distinct molecular signatures and phenotypic changes in cisplatin resistant PDX4 compared to their sensitive counterparts. Furthermore, we observed that chemoresistance was not inherently linked with increased stemness. In fact, although resistant cells expressed a combination of EMT and stemness markers, functional assays revealed that they were less proliferative, migratory, and clonogenic. JAK-STAT, hypoxia, and PI3K signaling pathways were enriched in these cells, indicating the activation of pathways that assist in DNA damage tolerance and cellular stress management.

**Conclusion:** This novel, syngeneic model provides a valuable platform for investigating the underlying mechanisms of cisplatin resistance in a clinically relevant context, contributing to the development of targeted therapeutic strategies tailored to combat resistance in stem-like ovarian cancer.

## 1 INTRODUCTION

In the United States, ovarian cancer ranks fifth among the leading causes of cancer death in women, making it the most lethal among gynecologic cancers^1^. Its lethality can often be attributed to the lack of effective screening for early detection^2^ and the recurrence that originates from chemotherapy resistant cells that persist through initial treatment^3^. Furthermore, ovarian cancer has been characterized as a largely heterogenous disease, resulting in varying effectiveness of currently available therapies^4,5^. Previously, our lab has focused on the characterization of a variety of patient-derived high-grade serous ovarian carcinoma (HGSOC) samples^6,7^. As the most aggressive subtype of ovarian cancer, the HGSOC samples indicated a large distribution of growth rate, morphology, gene and protein expression, and response to commonly used therapies, like cisplatin, a platinum-based agent that forms adducts with DNA, and olaparib, a poly (ADP-ribose) polymerase inhibitor (PARPi)^6^.

Multiple mechanisms of cisplatin resistance have been identified and extensively studied, including the regulation of drug influx/efflux, apoptosis, cell cycle, DNA repair pathways, among others^3^. Particularly in ovarian cancer, epithelial-mesenchymal transition (EMT), a dynamic and reversible biological process characterized by molecular and functional changes occurring in epithelial cells that result in a decrease in cell adhesion and increase in motility, has been linked to therapy resistance^8-11^. In this context, as cells transition to a more mesenchymal phenotype, chemoresistance has been correlated with the generation of cancer stem cells (CSC), which are a small subpopulation within tumors that are defined by their capacity for self-renew and differentiation^8,12-14^.

To better understand the development of chemoresistance and the potential novel mechanisms that could be utilized to target the disease, we sought to create a syngeneic model of cisplatin resistance from one of the patient-derived samples that was previously characterized, PDX4. Current available models of sensitivity and resistance in ovarian cancer are limited and do not depict the heterogeneity that is observed clinically^15,16^. Thus, our goal was to establish a methodology for the development of chemoresistance *in vitro* that parallel clinically relevant treatment concentrations. In this paper, we present the protocol utilized and characterize the cisplatin resistant PDX4 and its potential mechanisms for thwarting cell death.

## 2 RESULTS

### 2.1 Generation of cisplatin resistant cell line

PDX4, a high grade serous ovarian cancer (HGSOC) sample, was obtained as previously described from a chemotherapy naïve patient^6^. *In* vitro, the cells were described as epithelial-like, with low migration rate but high spheroid forming capacity when compared to additional patient-derived samples; this and other characteristics (including tumorigenicity) made this a sample with high stemness attributes ^6^. PDX4 was also determined to be *BRCA2* mutant and homologous recombination deficient by clinical testing^6^. Furthermore, *in vivo*, PDX4 demonstrated high tumorigenicity when orthotopically injected into the ovarian bursae of nude mice^6^. To further characterize this cell line, cisplatin resistant PDX4 (referred to as PDX4 CR throughout the remainder of paper) was generated through continual and repeated exposure to increasing concentrations of cisplatin (Figure 1A). A final cisplatin concentration of 8.5-10 µM was determined to be the maximal clinically relevant concentration that is observed in the plasma of patients; therefore, it was designated as the endpoint for further characterization^17-19^.

**Figure 1.**
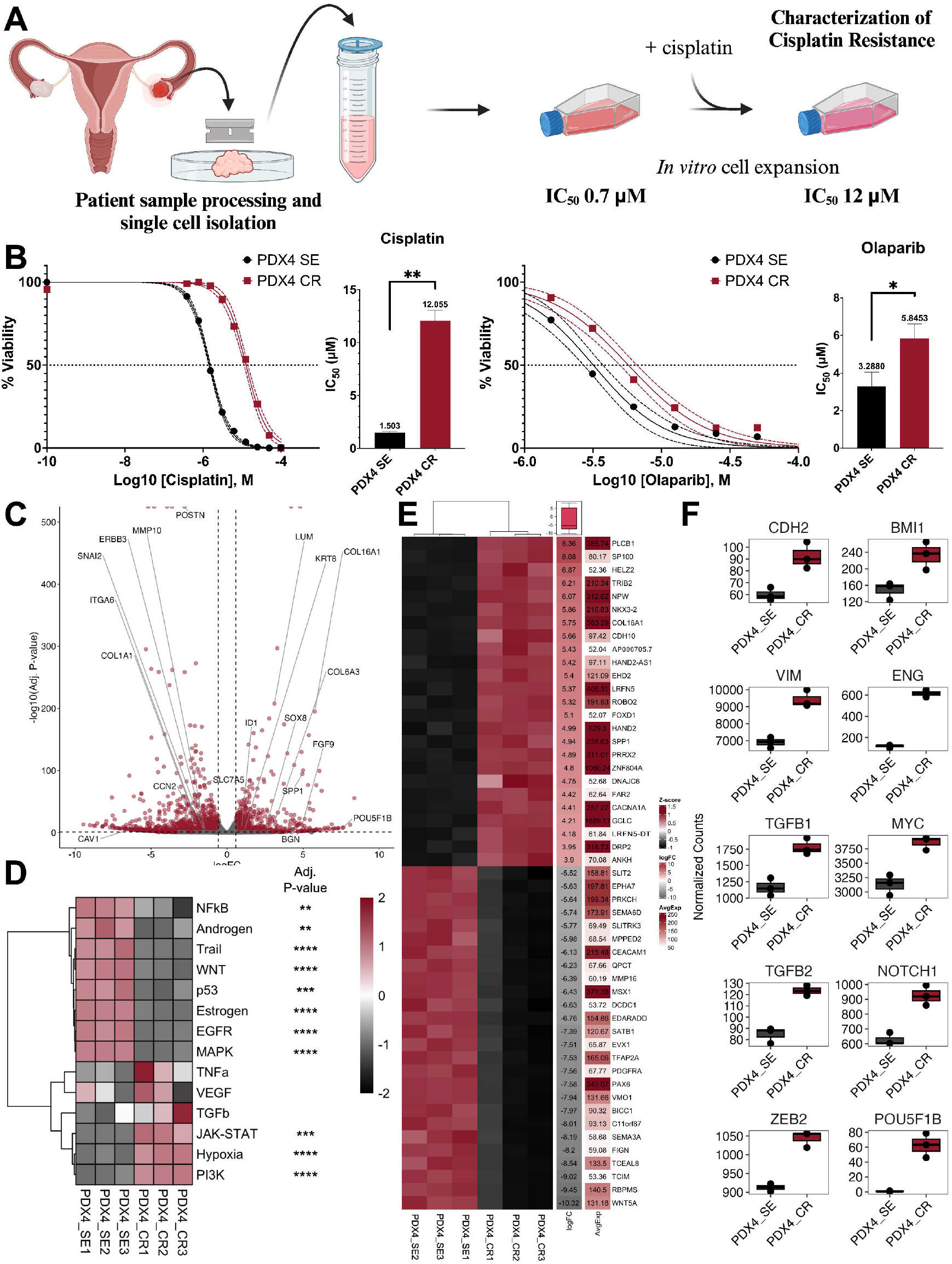
Cisplatin and olaparib resistant patient-derived cell line expresses markers of epithelial-mesenchymal transition (EMT) and stemness. A. PDX4 was obtained from a patient diagnosed with stage IVB high grade serous ovarian cancer (HGSOC). After sample dissociation and processing, cells were grown in vitro in increasing concentrations of cisplatin. B. While growing at 10 µM of cisplatin, PDX4 CR (cisplatin resistant) had cisplatin and olaparib IC50 of 12 µM (n=4) and 5.8 µM (n=3), respectively, as measured by thiazolyl blue tetrazolium bromide (methylthiazolyldiphenyl-tetrazolium bromide; MTT) viability assay. C. RNA sequencing revealed a total of 7,506 differentially expressed genes (DEGs), with 3,890 genes downregulated and 3,616 genes upregulated in PDX4 CR (n=3) relative to PDX4 SE (cisplatin sensitive; n=3). Each dot represents a single gene. The highlighted genes are associated with cisplatin resistance, cancer stemness, and/or EMT. Adjusted p-value < 0.1 and absolute log2(FC) > 0.58. D. PROGENy pathway signatures identified in cisplatin sensitive and resistant samples. E. Clustering heatmap displaying the top 50 differentially expressed genes (DEG) within the RNA-seq data. The logFC column indicates the expression relative to the cisplatin sensitive sample and the “AvgExp” column displays the average normalized count values across all samples. Above the logFC column, the box and whiskers plot summarizes the spread of the logFC expression values. F. Gene expression profiles of putative EMT (CDH2, VIM, TGFB1, TGFB2, and ZEB2) and stemness genes (BMI1, ENG, MYC, NOTCH1, and POU5F1B) in the cisplatin sensitive and cisplatin resistant samples as identified by RNA-seq. ^*^p < 0.05, **p < 0.01, ^***^p < 0.001. Created with Biorender.com.

Compared to vehicle treated control cells (cisplatin sensitive PDX4; PDX4 SE), PDX4 CR had an 8- and 1.8-fold increase in cisplatin and olaparib IC_50_, respectively (Figure 1B). At the transcript level, RNA sequencing revealed over 7,500 genes that were differentially expressed, with 3,890 genes that were downregulated and 3,616 genes that were upregulated in the chemoresistant cell line (adjusted p-value < 0.1; Figure 1C and Supplementary Table 2).

PROGENy algorithms were then utilized to identify signaling pathways that could be contributing to the differences observed between the two groups (Figure 1D). PDX4 CR displayed a significant activation of genes within the hypoxia, PI3K, and JAK-STAT signaling pathways, while NF-κB, androgen, TRAIL, WNT, p53, estrogen, EGFR, and MAPK pathways were significantly downregulated. Database searches revealed that, out of the top 50 differentially expressed genes, two genes have previously been associated with cisplatin resistance (*XAF1* and *TRIB2*)^20^, seven with EMT (*CCAT2, EVPL, SATB1, PDGFRA, BICC1, SFRP1, WNT5A*)^21^, and twenty-three with stemness^22^ (Figure 1E and Supplementary Table 3. Furthermore, markers and promoters of EMT, *CDH2, VIM, TGFB1, TGFB2*, and *ZEB2* were upregulated in the resistant sample, while genes associated with pluripotency and stemness, *BMI1, ENG, MYC, NOTCH1*, and *POU5F1B*, were upregulated in PDX4 CR (Figure 1F).

### 2.2 Growth and proliferation characteristics after the acquisition of cisplatin resistance

To verify cell viability, PDX4 SE and CR were treated with cisplatin for 72 hours, imaged, then labeled with Annexin V and 7AAD and quantified with flow cytometry (Figure 2A-B). After cisplatin treatment, under phase-contrast micrographs, PDX4 SE displayed mild morphological changes, acquiring a more elongated shape (Figure 2A; left). In contrast, PDX4 CR maintained its epithelial-like, cobblestone pattern even at high concentrations of cisplatin (Figure 2A; right). Furthermore, PDX4 SE had a large decrease in cell confluency with the first concentration of cisplatin (3.125 μM), while PDX4 CR gradually decreased in confluency (Figure 2A). More specifically, at 3.125 μM cisplatin, PDX4 SE had around 2-4% live cells (Annexin V^-^/7AAD^-^; Figure 2B, top), 30-59% cells undergoing early apoptosis (Annexin V^+^/7AAD^-^; Figure 2B, middle), and 32-54% cells in late apoptosis and necrosis (Annexin V^+^/7AAD^+^; Figure 2B bottom). In comparison, the percentage of live cells for PDX4 CR only decreased considerably from 6.25 μM (42-84%) to 12.5 μM (12-51%) of cisplatin (Figure 2B). At 12.5 μM and 50 μM of cisplatin, PDX4 CR had the highest percentage of early apoptotic (51%) and late apoptotic/necrotic cells (75%), respectively (Figure 2B).

**Figure 2.**
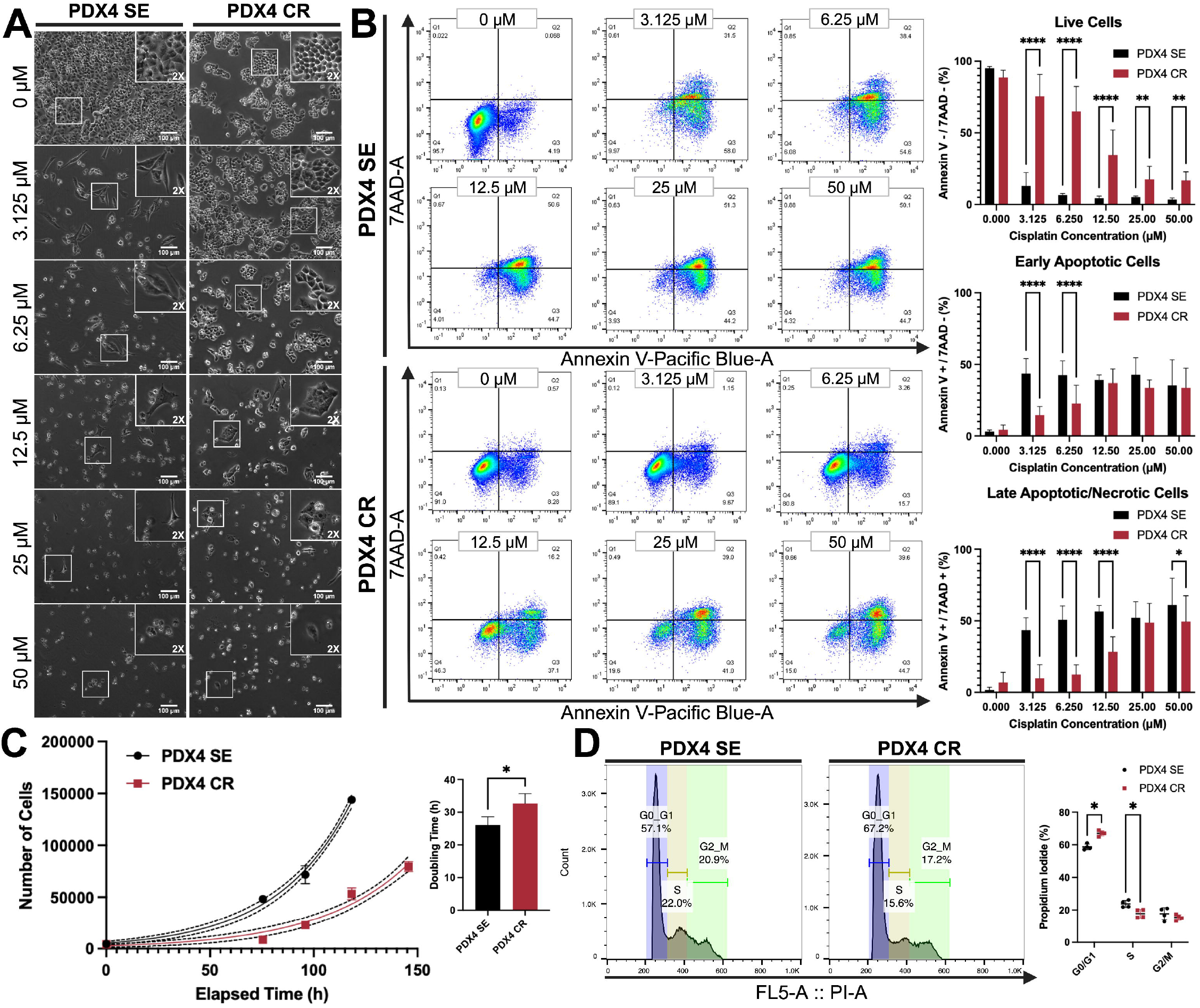
Growth characteristics of PDX4 CR after cisplatin treatment. A. After three days of cisplatin treatment (0 – 50 µM), PDX4 SE and CR were imaged at 10X magnification. Cell morphologies are highlighted in each panel with an inset at 2X magnification. Scale bar: 100 µm. B. Representative FACS profiles of PDX4 SE and CR cells stained with Annexin V-Pacific Blue (PB) and 7-aminoactinomycin D (7AAD) after 72 h of cisplatin treatment (0 -50 µM). Quantification of live Annexin V-/7AAD-(top), early apoptotic Annexin V+/7AAD-(middle), and late apoptotic/necrotic Annexin V+/7AAD+ and Annexin V-/7AAD+ (bottom) cells. Percent positive cells were gated from single, intact cells. C. Analysis of cell proliferation by MTT assay. PDX4 SE and CR were seeded at 1,000 cells per well on day zero and allowed to grow for 96 – 144 h (4 – 6 days). Doubling time was determined from exponential (Malthusian) growth curve (n = 3). Data is presented as mean ± SD. Curve fitting and doubling time were calculated with GraphPad Prism software. D. Cell cycle analysis by flow cytometry (left). Untreated PDX4 SE and CR cells were stained with FxCycle™ propidium iodide/RNase staining solution. Percent distribution of cells through the different cell cycle stages (right). All flow cytometry data was collected with a MACSQuant Analyzer 10 and analyzed with FlowJo software. Created with Biorender.com.

As cisplatin sensitive and resistant cells were grown in parallel *in vitro*, we observed that PDX4 CR seemed to proliferate slower. In fact, quantification of doubling time revealed that PDX4 SE had an average doubling rate of 26 hours, while PDX4 CR doubled every 33 hours (Figure 2C). Additionally, both cell lines were stained with propidium iodide (PI) for cell cycle analysis through flow cytometry (Figure 2D). Interestingly, PDX4 CR had a higher percentage (67%) of cells in the G0/G1 phase than PDX4 SE (59%; Figure 2D), indicating that the cells were likely less proliferative due to a portion of them being in cell cycle arrest. At the S phase, a slightly higher proportion of sensitive cells (24%) than resistant cells (18%) was found.

### 2.3 Defining Epithelial Mesenchymal Transition Status

Previous literature indicates that platinum-based therapies may induce the epithelial-mesenchymal transition (EMT) process^23-27^. Since the RNA-seq data highlighted pathways activated in EMT (Figure 1D) along with the differential expression of several EMT markers (Figure 1F), we set out to validate not only the EMT-associated phenotypical and functional changes observed in PDX4 SE and CR, but also the mechanism of cisplatin resistance.

We used RT-qPCR to validate the mRNA-seq results. In mRNA expression, PDX4 CR had a significant increase in the expression of *SNAI1, ZEB2*, E-cadherin (*CDH1*), and N-cadherin (*CDH2*), while PDX4 SE had a higher expression of Slug (*SNAI2*) (Figure 3A). Interestingly, at the protein level, Snail expression was only slightly increased, and vimentin was decreased significantly in PDX4 CR (Figure 3B and Supplementary Figure 3). Additionally, flow cytometry identification of E-cadherin (CD324) and N-cadherin (CD325) revealed that PDX4 CR had more CD324^+^ (50%) and CD325^+^ (35%) cells than the cisplatin resistant cells (41% and 26%, respectively; Figure 3C).

**Figure 3.**
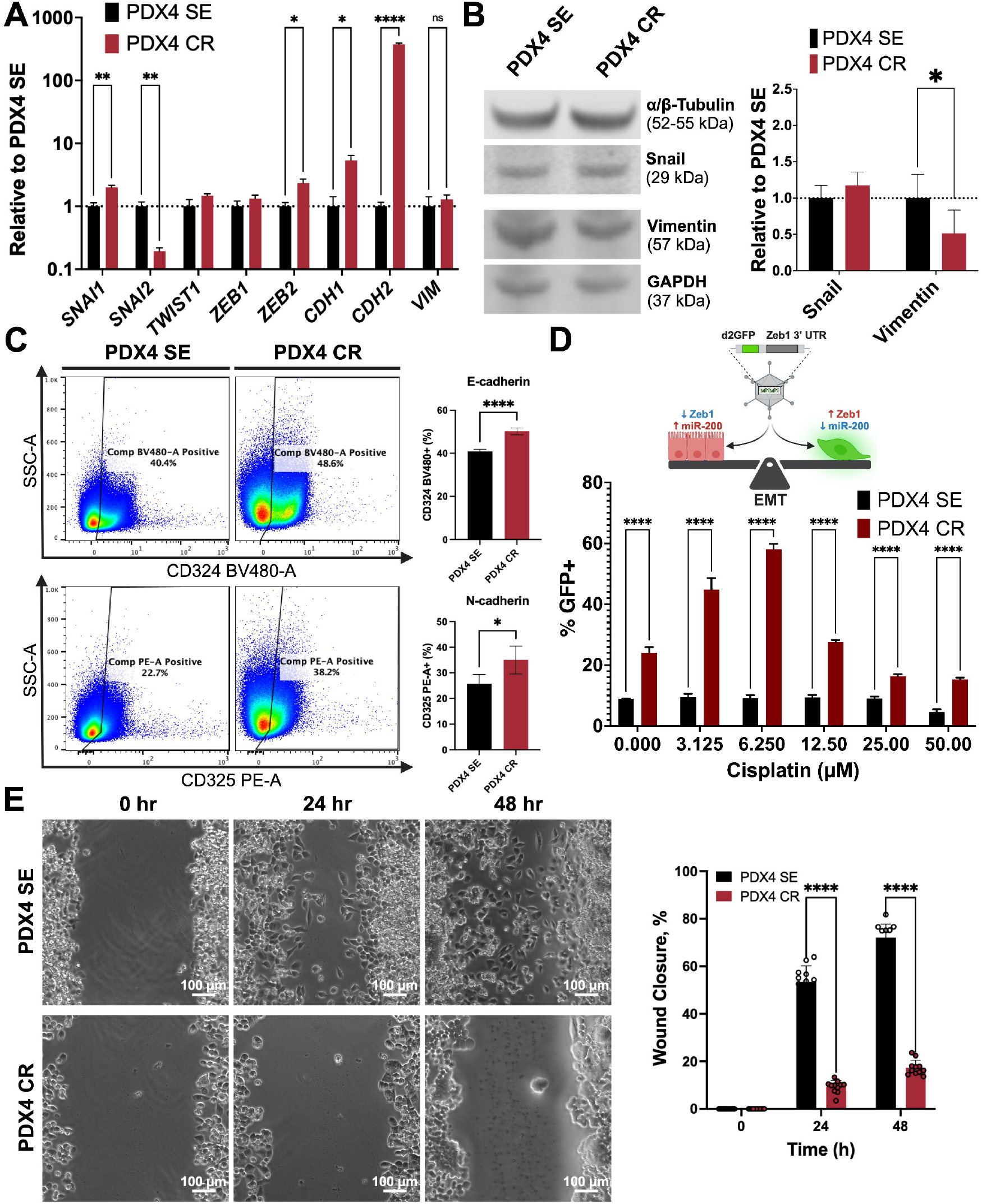
PDX4 SE and CR cells exhibit hybrid/partial EMT phenotype. A. RT-qPCR validation of EMT transcription factors and markers. Expression levels are relative to the sensitive cell line. Results are presented as the means ± SD. B. Protein quantification of Snail and Vimentin through western blot, normalized to either α/β-tubulin or GAPDH. Expression levels are relative to PDX4 SE. C. Representative flow cytometry dot plots (left) for the expression of E-cadherin (CD324) and N-cadherin (CD325). Quantification of single and double positive populations (right). Percent positive cells were obtained from single, intact, live cells. D. The Zeb1 3’ UTR GFP reporter (Toneff et al., 2016) detects the presence of miR-200 within cells. PDX4 SE and CR cells were transduced with the lentiviral reporter system and treated with cisplatin (0 -50 µM) for 72 h. Quantification of GFP expression was measured through flow cytometry. GFP positive populations were obtained from single, intact cells that were Annexin V-/7AAD-. All flow cytometry data was collected with a MACSQuant Analyzer 10 and analyzed with FlowJo software. E. Migration capacity was determined through 24- and 48-h scratch wound healing assays (n=12). Results are presented as the means ± SD. Statistical significance was determined by two-way ANOVA with Tukey’s test for multiple comparison correction. P-values: P ≤ 0.05 (^*^); P ≤ 0.01(**); P ≤ 0.001 (^***^); P ≤ 0.0001 (^****^). Created with Biorender.com.

Since EMT is a transitional process, we aimed to describe the effect of different concentrations of cisplatin on the expression of EMT markers on a per cell basis. For this purpose, we transduced PDX4 SE and CR with the *ZEB1* 3’ UTR destabilized GFP (dGFP) reporter^28^ that senses EMT changes as a function of miR-200 levels. More specifically, as Zeb1 levels increase with EMT, miR-200 repression allows dGFP expression, detected through flow cytometry (Figure 3D). For all concentrations tested, GFP positive populations were greater in the chemoresistant cells, with the highest enrichment achieved at 6.25 μM.

Functionally, migratory capacity was measured through a wound healing assay. Because of gene expression changes consistent with a more mesenchymal phenotype overall, it was expected that PDX4 CR would be more migratory. However, after scratching the monolayer of cells, PDX4 SE displayed a higher efficiency in gap closure compared to PDX4 CR (Figure 3E). By 24 hours, PDX4 SE cells at the leading edge spread towards the center of the gap, while PDX4 CR cells appeared stationary, with their leading edge mainly intact (Figure 3E).

### 2.4 Stemness and Self-renewal Characteristics

Stemness, within the context of cancer, refers to the ability of cancer cells to self-renew and differentiate into heterogeneous populations within a given tumor. In many cancer types, activation of EMT has often been associated with the generation of stemness characteristics^12,13,29^. Given that both the cisplatin sensitive and resistant cells expressed EMT markers, we sought to characterize their stemness potential.

PDX4 SE had a higher expression of *LIN28A*/Lin28a at both the mRNA and protein level than its resistant pair, while PDX4 CR expressed more *NANOG* and Oct4 at mRNA and protein level, respectively, relative to PDX4 SE (Figure 4A-B). Corroborating the LIN28 expression patterns, we observed an overall enrichment in the expression of let-7 family members in the resistant cells, microRNAs known to promote a differentiated state, acting in a negative feedback loop in the regulation of LIN28 expression (Supplementary Figure 4 and Supplementary Table 4). Furthermore, flow cytometry was used for the quantification of cells positive for CD44, CD117, CD133, and aldehyde dehydrogenase (ALDH) activity, all markers of stemness that have been validated in ovarian cancer^30,31^. All tested markers were expressed abundantly in both sensitive and resistant cells (Figure 3C), but PDX4 CR had a significantly higher proportion of all markers. Specifically, more than half of the live population of cells were CD44^+^ in both PDX4 SE (54%) and PDX4 CR (67%; Figure 3C). Likewise, CD117^+^, CD133^+^, and ALDH^+^ populations were 2.5-, 1.2-, and 1.6-fold greater, respectively, in the resistant cells compared to their sensitive counterpart (Figure 3C).

**Figure 4.**
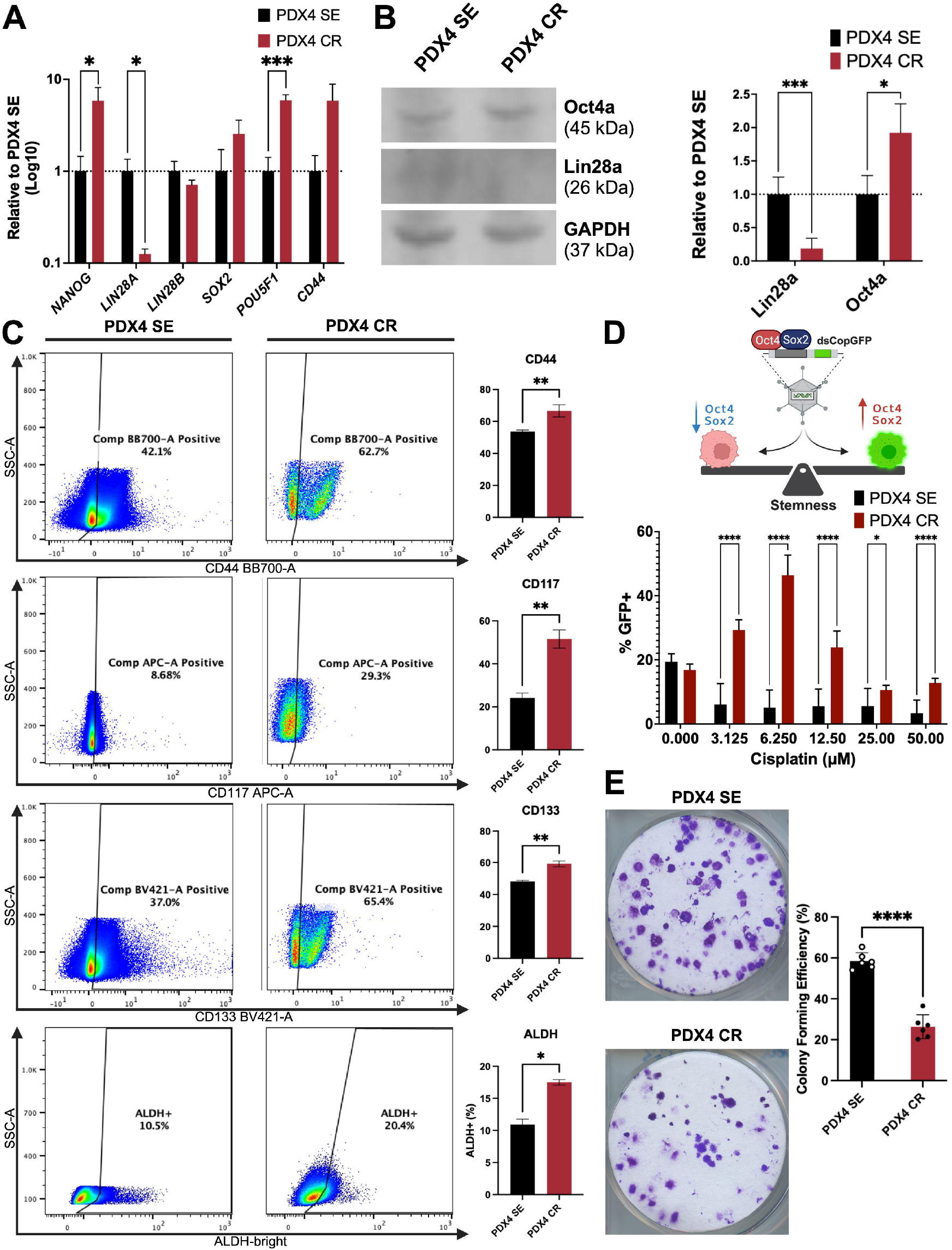
Cisplatin resistant cells express more stemness markers but are less clonogenic. A. RT-qPCR validation of pluripotency and stemness markers. Expression levels are relative to the sensitive cell line. Results are presented as the means ± SD. B. Protein quantification of Lin28a and Oct4a through Western blot, normalized to GAPDH, relative to PDX4 SE. C. Representative flow cytometry dot plots (left) for the expression of cell surface markers CD44 (first row), CD117 (c-Kit; second row), CD133 (PROM1; third row), and ALDH (last row). Comparison of single positive populations between PDX4 SE and CR (right). Percent positive cells were obtained from single, intact cells. ALDH gates were drawn based on DEAB treated cells. D. The SORE6 GFP reporter (Tang et al., 2015) detects the presence of Sox2 and Oct4 within cells. PDX4 SE and CR cells were transduced with the reporter system and treated with cisplatin in increasing concentrations (0 - 50 µM) for 72 h. Quantification of GFP expression was measured through flow cytometry. GFP positive populations were obtained from single, intact cells that were Annexin V-/7AAD-. All flow cytometry data were collected with a MACSQuant Analyzer 10 and analyzed with FlowJo software. E. Representative images of PDX4 SE and CR colonies that were grown for 10-11 days (left). Images were acquired with an HP scanner. Colony forming efficiency was calculated relative to the number of plated cells (n=6; right). Results are presented as the means ± SD. Statistical significance was determined by unpaired T-test. P-values: P ≤ 0.05 (^*^); P ≤ 0.01(**); P ≤ 0.001 (^***^); P ≤ 0.0001 (^****^). Created with Biorender.com.

Taking into consideration that stem cells are a rare population, we aimed to describe the effect of different concentrations of cisplatin on the expression of pluripotency markers at a per cell basis. Thus, we transduced PDX4 cells with the SORE6-GFP reporter^32^, with GFP expression under the control of the Sox2/Oct4 response element (SORE). Notably, pluripotent cells would express more Sox2 and Oct4, resulting in an increase in the translation of GFP that can be detected through flow cytometry (Figure 3D). For all concentrations tested (including in the untreated cells), GFP expression was greater in the chemoresistant cells, with the highest enrichment achieved at 6.25 μM (Figure 3D).

To test self-renewal capacity, sensitive and resistant PDX4 cells were subjected to colony forming assays. As a result, PDX4 SE demonstrated higher efficiency at colony formation with an average of almost 60% of cells plated being able to form colonies. In comparison, around 25% of PDX4 CR cells demonstrated colony forming ability (Figure 3E). Together, these results indicate that, while PDX4 SE and CR had different stem-like expression patterns, PDX4 SE had higher self-renewal capacity.

## 3 DISCUSSION

Many mechanisms of cisplatin resistance have been described in the literature^3^. In accordance with previous findings, we had hypothesized that resistance would correlate with a more mesenchymal-like status and increased stemness^23-27^; however, we observed that, although cisplatin resistant cells retained expression of EMT and stemness markers, functional findings were not in agreement. Furthermore, with the RNA-seq results, PDX4 CR had an enrichment in the expression of genes associated with the hypoxia, PI3K/Akt, and JAK-STAT pathways, all of which have been associated with EMT to different extents^33-37^. Interestingly, PDX4 SE and CR were both classified as Stem-A, with the sensitive cell line showing greater enrichment of Stem-A signature compared to the resistant cells (Supplementary Figure 5). The Stem-A ovarian cancer subtype has been described as corresponding to the Tothill et al. C5 and the TCGA proliferative subtypes^4^. In terms of gene expression profile, this subtype is known for demonstrating an enrichment in the expression of genes associated with development, proliferation, and stemness^4^. Moreover, its aggressiveness in ovarian cancer has been attributed to the activation of non-canonical Wnt/PCP pathway through *FZD7*^38^. Gene set enrichment analysis confirmed that the non-canonical Wnt pathway was activated in PDX4 SE; however, PDX4 CR did not exhibit such patterns (Figure 1, Supplementary Figure 6). In fact, the cisplatin resistant cells demonstrated an enrichment in pathways associated with immune signaling, protein localization, and mitochondrial function (Supplementary Table 5 and 6). The shift away from Stem-A could potentially be explained by PDX4 CR cells utilizing these enriched signaling cascades to hijack pathways aiding DNA damage tolerance. Further studies are underway to elucidate such novel mechanisms contributing to resistance and the shift away from the Stem-A/C5/Proliferative subtype.

In addition to the analysis of pathways, we performed an initial assessment of multiple cisplatin and copper transporters through qPCR and flow cytometry; however, no significant difference was detected between the cisplatin sensitive and resistant cells (Supplementary Figure 7). Furthermore, while testing DNA damage repair capabilities, western blot quantification of PARP1 indicated that PDX4 CR had lower expression of the protein (Supplementary Figure 3C). Considering that PDX4 samples are BRCA2 mutant, treatment with olaparib should have induced synthetic lethality; however, the resistant cells had increased resistance to the PARPi, suggesting the potential for higher DNA damage repair proficiency. As a response to cisplatin treatment (presumably the DNA damage caused by that drug), we observed that cells proliferated at a slower pace (Figure 2C), remaining in the earlier stages of the cell cycle, G0/G1 (Figure 2D). The delay in proliferation could be the explanation of why the sensitive cells appear to be more migratory and clonogenic. Future studies should focus on determining whether cisplatin withdrawal would result in the restoration or improvement of these cellular functions.

Alternatively, the accumulation of cells in G0 could indicate that the resistant cells are transitioning into a state of dormancy while facing genomic instability or biological stress. This interpretation would agree with the assessment that PDX4 CR cells may be activating the cGAS-STING signaling pathway, which has been associated with cell cycle arrest^39-43^.

Taking together the complex nature of these results and the mechanism of action of cisplatin, we reason that PDX4 CR cells maintain and lose some EMT- and stem-like characteristics to undergo functional changes towards dormancy for the promotion of DNA damage repair and, thereby, survival. With this study, we have developed a novel syngeneic model of cisplatin resistance in which to study the effect of treatment on stem-like ovarian cancer cells. To our knowledge, this model is the first to demonstrate the aforementioned mechanism of cisplatin resistance within ovarian cancer. Additionally, with this model, there is the potential to better understand the interface between DNA damage and immune response and to determine if this pathway is targetable for the enhancement of patient response to immunotherapies.

## 4 METHODS

### 4.1 Cell Culture

PDX4, a chemotherapy naïve HGSOC sample, was obtained as previously described ^6^. Briefly, with informed consent, tumor tissue was collected by the Loma Linda University Cancer Center Biospecimen Laboratory (LLUCCBL). The same tumor samples were used to clinically diagnose HGSOC. To obtain a single cell suspension, the tissue section was washed in 1X phosphate-buffered saline (PBS; 138 mM NaCl, 2.7 mM NaCl, 8.1 mM Na_2_HPO_4_, 1.2 mM KH_2_PO_4_ in ddH_2_O) containing 20 μg/ml gentamicin sulfate (G020-1GM; Caisson Labs, Smithfieeld, UT, USA) minced with a razor blade (SKU 71964; Electron Microscopy Sciences, Hatfield, PA, USA), and passed through a 100-μm cell strainer (15-1100; Biologix, Camarillo, CA, USA) using a cell pestle (229480; CELLTREAT Scientific Products, Pepperell, MA, USA). Erythrocytes were removed with Cytiva Ficoll-Paque™ PLUS media (17144002; Cytiva, Marlborough, MA, USA) centrifugation. Cells were cultured long-term in a 3:1 mixture of Hyclone™ Ham’s Nutrient Mixture F12 with L-glutamine (SH30026.01; Cytiva) and Dulbecco’s Modified Eagle’s Medium with high glucose and L-glutamine (DMEM; 25-501; Genesee Scientific, El Cajon, CA, USA), supplemented with 5% FBS (Omega Scientific, Tarzana, CA, USA), 0.4 μg/ml hydrocortisone (H0888-1G; Sigma-Aldrich, St. Louis, MO, USA), 5 μg/ml insulin (91077C-100MG; Sigma-Aldrich), 2 μg/ml isoprenaline hydrochloride (I5627-5G; Sigma-Aldrich), 24 μg/ml adenine (A8626; Sigma-Aldrich), 100 U penicillin, and 100 μg/ml streptomycin (25-512; Genesee Scientific). Cells were maintained at 37°C with 5% CO_2_ and tested for mycoplasma contamination routinely, using the PlasmoTestTM Mycoplasma Detection Kit (rep-pt1; InvivoGen, San Diego, CA, USA).

### 4.2 Generation of Cisplatin Resistant Cell Line

PDX4 cells were plated at 25% confluence in three 25 cm^2^ tissue culture flasks (12-556-009; Thermo Fisher Scientific, Waltham, WA, USA). After 24 hours, one flask received vehicle treatment (PBS), a second flask received 0.7 μM cis-Diammineplatinum(II) Dichloride (cisplatin; IC-50 determined previously ^6^; D3371-100MG; TCI Chemicals, Portland, OR, USA), and the last flask was used as a plating, health assessment control. Once cells achieved 70-80% confluence, they were passaged three more times within the same concentration of cisplatin before treatment concentration was increased by 0.05, 0.1, and then finally 0.5 μM increments until they were able to withstand and grow healthily at 10 μM cisplatin, which was determined to be the maximal plasma concentration observed in patients ^17-19^.

One millimolar cisplatin aliquots were prepared in large batches and frozen at -70°C to prevent batch variation and degradation through multiple freeze-thaw cycles.

### 4.3 Cell viability assays

Cell viability was measured using thiazolyl blue tetrazolium bromide (MTT; 00697; Chem-Impex, Wood Dale, IL, USA) assays. One thousand PDX4 cells were seeded in triplicates in flat-bottomed 96-well plates (229195; CELLTREAT Scientific Products) and allowed to adhere for 24 hours prior to the addition of cisplatin or Olaparib (AZD2281; S1060; Selleck Chemicals, Houston, TX, USA). Throughout the duration of the experiments, growth medium without penicillin-streptomycin was used. Cells were then treated with vehicle (PBS for cisplatin and DMSO for olaparib) or increasing concentrations of cisplatin or olaparib (0-100 μM) followed by an incubation of 72 hours. At the endpoint, plates were treated with 5 mg/ml MTT (solubilized in PBS) and incubated for 3 hours at 37°C. After confirmation of formazan crystal formation, wells were aspirated before the addition of 100 μL dimethyl sulfoxide (DMSO; 0219605591; MP Biomedicals, Santa Ana, CA, USA) per well. Absorbance was measured at 560 nm using a SpectraMax i3x microplate reader (Molecular Devices LLC, San Jose, CA, USA).

Absorbance readings were normalized to vehicle treated values, and the half-maximal inhibitory concentration (IC-50) of drug was determined in cisplatin sensitive and resistant cells using GraphPad Prism v 9.5.0 (GraphPad Software, La Jolla, CA, USA).

### 4.4 Library Preparation and RNA-sequencing

Total RNA was extracted from PDX4 SE and PDX4 CR cells using the miRNeasy Mini Kit (217004, Qiagen). Before sequencing, RNA quality was checked through agarose gel electrophoresis using 2X RNA loading dye (R0641; Thermo Fisher Scientific) according to the manufacturer’s protocol. All RNA samples were derived from three independent experiments. Subsequently, RNA-seq library construction and generation of raw data was performed at the Loma Linda University Center for Genomics. RNA-seq libraries were constructed using the Ovation Universal RNA-seq System (0364; Tecan; Männedorf, Switzerland). Briefly, 100 ng of total RNA was reverse transcribed and then made into double stranded cDNA by the addition of a DNA polymerase. cDNA was concentrated using Agencourt beads followed by end repair and adaptor ligation. Unique barcodes were used for each sample for multiplexing. Targeted rRNA-depletion was performed before final library construction. Libraries were amplified using 13 cycles in the Eppendorf™ Mastercycler™ pro PCR system (Hamburg, Germany) and purified using Agencourt beads. Library QC was performed with a TapeStation 2200 (Agilent Technologies) and all libraries were quantified using Qubit 4.0 (Life Technologies). RNA-seq libraries were sequenced on Illumina NextSeq 550 (Illumina, San Diego, CA, USA) with single 76 bp reads. Illumina RTA v2.4.11 software was used for basecalling and bcl2fastq v2.17.14 was used for generating FASTQ files.

### 4.5 Bioinformatics Analysis

Sample quality was determined with FASTQC (v 0.11.9) ^44^ and MultiQC (v 1.11) before and after reads were trimmed with Trimmomatic (v 0.39) ^45^. Subsequently, reads were aligned to the human genome with STAR (v2.7.10a) ^46^, using GRCh38 ^47^, release 109, as reference. BAM (binary alignment map) files that were generated from STAR alignment were sorted and indexed with SAMtools (v 1.12) ^48^. Further processing of reads involved: 1) utilizing HTSeq (v 2.0.2) for the assembly of RNA-seq reads into transcripts; 2) counting transcripts; 3) producing a count matrix from the HTSeq output for the differential gene expression analysis between samples with DESeq2 (v 1.43.1) ^49,50^. To enhance the robustness of gene expression analysis, genes with less than 10 counts in all samples were excluded from the matrix. Downstream analysis required the conversion of Ensembl identifiers (ID) to HGNC symbols, which was done with the BioMart R package (v 2.58.0) ^51^.

### 4.6 Pathway analysis

The PROGENy (v 1.24.0) ^52^ R package was used for inference of pathway activity. DEGs were selected based on whether their expression had an adjusted p-value < 0.1 and absolute FC greater than 1.5. Genes that did not fit these criteria were filtered out from the normalized count matrix, which was used as input for PROGENy. Cisplatin resistant samples were compared to sensitive samples with a linear regression model. The p-values obtained were adjusted using the Banjamini-Hoechberg false discovery rate (FDR) method. Activity was considered significant if FDR was < 0.1.

### 4.7 Microscope Imaging

Cell imaging acquisitions were done with a Nikon Eclipse Ti microscope and μManager v 1.4.22 software. Unless otherwise noted, images were taken at 10X magnification.

Magnification insets were generated and added to image panels with the QuickFigures plugin in ImageJ.

### 4.8 Flow Cytometry

The apoptosis assay was conducted using Annexin V-Pacific Blue (640918; Biolegend, San Diego, CA, USA) and 7AAD (13-6993-T500; Tonbo Biosciences, San Diego, CA, USA). One hundred thousand cells were plated per well in a 12-well plate. Cells were treated with cisplatin for 72 hours and then harvested in PBS. After centrifugation, cells were resuspended in 5 μL of Annexin V and 10 μL Annexin V binding buffer (20 mM HEPES, 140 mM NaCl, and 2.5 mM CaCl_2_ in ddH_2_0) and incubated for 15 min at 4°C. Next, cells are stained with 5 μL of 7AAD for 5 min at 4°C. Once incubation period ends, cell suspension is diluted with 180 μL of Annexin V binding buffer. Single stain controls were also plated for both Annexin V-PB and 7AAD.

For cell cycle analysis, PDX4 CR cells were cultured without Cisplatin (10uM) for 1 week and collected before assay was performed. PDX4 SE and CR were collected via trypsinization, counted to 1x10^6^, rinsed with room temperature 1x sterile PBS twice, fixed with chilled 70% EtOH, and rinsed with 1x sterile PBS twice. Cells were resuspended in 500 μL of FxCycle™PI/RNase Staining Solution (F10797, lot# 2561624, Invitrogen, Oregon) and left to incubate in the dark for 30 minutes.

Cells were resuspended in FACS stain [1% FBS, 0.1% sodium azide (NaN_3_; s2002-5g; Sigma), and 2 mM EDTA in PBS] or BD Horizon Brilliant Stain Buffer (563794; BD, Franklin Lakes, NJ, USA), and labeled with conjugated fluorescent dye antibodies against CD44 BB700 (745949), CD117 APC (567128), CD133 BV421 (566598), CD324 BV480 (E-Cadherin;

752951), and CD325 PE (N-Cadherin; 561554) obtained from BD. After incubation at 4°C for 30 min, cells were washed and resuspended in FACS stain. UltraComp eBeads (01-2222; Thermo Fisher Scientific) and BD Horizon™ fixable viability stain 780 (565388) were used for compensation and live cell gating, respectively. ALDH positive cells were identified with the ALDEFLUOR™ assay kit (01700; STEMCELL Technologies, Vancouver, Canada), using the manufacturer’s instructions. Briefly, one million cells were resuspended in 1mL of ALDEFLUOR™ Assay Buffer. Subsequently, five microliters of activated ALDEFLUOR Reagent were added to the cell suspension and mixed. Half of the cell suspension was immediately transferred to the DEAB Reagent control tube before incubation for 30 min at 37°C.

Flow cytometry was performed on MACSQuant Analyzer 10 (Miltenyi Biotec, Bergisch Gladbach, Germany), and analysis of data was performed using FlowJo 10 (FlowJo LLC, Ashland, OR, USA).

### 4.9 Proliferation Assay

Cells were seeded in two flat-bottomed 96-well plates: the test plate was seeded with 1,000 cells/well and the standard growth curve plate with a two-fold serial dilution of cell suspension, ranging from 0 to 50,000 cells/well in triplicates. After incubation for 24 h, the standard growth curve plate and quadruplicates of the test plate were treated with 5 mg/ml MTT (solubilized in PBS) and incubated for 3 h at 37°C. After confirmation of formazan crystal formation, wells were aspirated before the addition of 100 μL DMSO per well. Absorbance was measured at 560 nm using a SpectraMax spectrophotometer. Absorbance readings from the standard growth curve were normalized to blank wells before being calculated into relative viable cell count. Remaining wells from the test plate were harvested in quadruplicates in the following days (up to 7 days) for the calculation of doubling time. After each reading, wells were washed at least three times once DMSO was removed.

### 4.10 Quantitative Reverse Transcription PCR (RT-qPCR)

Total RNA from cell culture samples was isolated using IBI Isolate DNA/RNA Reagent Kit (#IB47602, IBI Scientific, Dubuque, IA, USA) according to the manufacturer’s instructions. After RNA purification, cDNA was synthesized from 1 μg of total RNA using Maxima First Strand cDNA Synthesis Kit (K1672; Thermo Fisher Scientific). RT-qPCR was performed using Applied Biosystems™ PowerUP™ SYBR™ Green Master Mix (A25778; Thermo Fisher Scientific) and specific primers on a Stratagene Mx3005P Instrument (Agilent Technology, Santa Clara, CA, USA). The sequence of all human primers is shown in Supplementary Table 1. The results were analyzed using the ΔΔ cycles to threshold (ΔΔ*C*_t_) method^53^.

### 4.11 Protein Extraction and Quantification

To quantify protein expression, cisplatin sensitive and resistant cells were plated in triplicates in 6-well (100,000 cells/well) or 12-well plates (50,000 cells/well) and allowed to expand for 72 h before trypsinization, PBS washing, and addition of Laemmli sample buffer [0.5M Tris HCl at pH 6.8 (50-213-711; Fisher Scientific), 10% sodium dodecyl sulfate (SDS), 100mM PMSF (329-98-6; Sigma-Aldrich) in isopropanol (190764-4x4l; Sigma), glycerol (G5516-1L; Sigma-Aldrich), and protease inhibitor (11836170001; Sigma-Aldrich) dissolved in ddH_2_O]. Lysates of cells were sonicated with a Sonic Dismembrator Model F60 (remote, 10 seconds at 50% power, then 2 minutes on ice, repeated three times; Fisher Scientific) and passed through a Hamilton syringe (14-824-663; Thermo Fisher Scientific) to dissociate proteins attached to chromatin. Subsequently, samples were centrifuged at 10,000 rpm for 10 minutes and supernatant was transferred to a new tube. Quantification of proteins was carryout out with a BCA protein assay (23227; Thermo Fisher Scientific). Absorbance was measured at 562 nm using a SpectraMax i3x microplate reader (Molecular Devices LLC).

### 4.12 Electrophoresis and Immunoblotting

Loading buffer [8% SDS, 10% 2-mercaptoethanol (21985023; Life Technologies, Carlsbad, CA, USA), 30% glycerol, 0.008% bromophenol blue (BP115-25; FisherScientific, Fair Lawn, NJ), and 0.2 M Tris HCl in ddH_2_O] was added to 50 μg of proteins from whole cell lysates, then samples were heated to 100°C for 5 min before being loaded into individual lanes of 4-12% SurePAGE™ Bis-Tris polyacrylamide gels (GenScipt, Piscataway, NJ, USA) and separated with electrophoresis within MES SDS running buffer (M00677; GenScript), using a Mini PREOTEAN 3 cell (525BR058974; Bio-Rad, Hercules, CA, USA) and PowerPac™ Power Supply (043BR09142; Bio-Rad) at 150 V for 1 h and 45 min in a cold room. Proteins were then transferred to a 0.45 μm Immobilon-FL PVDF membranes (IPFL00010; EMD Millipore, Burlington, MA, USA) using the eBlot™ L1 Fast Wet Transfer System (GenScript) according to the manufacturer’s instructions. Membranes were dried at room temperature for 1 hr, re-activated with methanol (MX0485-7; EMD Millipore), and blocked with 5% milk in TBS [20 mM Tris– HCl, pH 7.6, and 140 mM NaCl (SB0476-1; Bio Basic, Markham, Ontario, Canada) in ddH_2_O] for 1 h. Membranes were probed with corresponding primary antibodies at a 1:1000 dilution (5% milk in TBST) overnight at 4°C. Primary antibodies (Cell Signaling Technologies, Danvers, MA, USA) included mouse anti-Snail (3895S), rabbit anti-Vimentin (5741), rabbit anti-LIN28A (3978), rabbit anti-OCT4A (2840), rabbit anti-α/β-tubulin (2148), and rabbit anti-GAPDH (2118). Secondary antibody immunoblotting was done with goat anti-rabbit IgG (IRDye 680RD; 926-68071; LI-COR Biosciences, Lincoln, NE, USA, or DyLight 680; PI35569; Thermo Fisher Scientific) and goat anti-mouse DyLight 800 (SA5-10176; Invitrogen, Waltham, MA, USA) for an hour at a 1:30,000 working concentration (5% milk in TBST with 0.01% SDS) at room temperature. Membranes were imaged with LI-COR Odyssey CLx Infrared Imaging System (LI-COR Biosciences) and analyzed with ImageJ software v1.54f (National Institutes of Health, Bethesda, MD, USA) and ImageStudio (LI-COR Biosciences). Full blots are shown in Supplementary Figure 1 and 2.

### 4.13 GFP reporters

FUGW-eGFP-ZEB1 3’UTR was a gift from Dr. Jeffrey Rosen (Addgene plasmid #81018; http://n2t.net/addgene:81018; RRID: Addgene_81018). SORE6-GFP lentiviral reporters were a gift from Dr. Lalage M. Wakefield. Both plasmids were selected with ampicillin in Stbl3 *E. coli* (C737303; Invitrogen) cultures and verified with DNA digestion and gel electrophoresis before clonal expansion and purification with QIAGEN Plasmid Maxi Kit was performed (12163; Qiagen, Redwood City, CA, USA). Lentivirus particles were produced in HEK293T cells after co-transfection of lentivirus plasmid vectors with packaging plasmids (VSVG and lenti gag/pol) using polyethylenimine (PEI; NC1014320; Polysciences, Warrington, PA, USA). After 48 h and 72 h, medium containing lentivirus was collected and filtered through 0.22 μM filter. Filtered virus containing medium was used for cell transduction or stored at −70 °C. Cells were transduced with lentivirus in the presence of 6 μg/mL protamine sulfate (p3369-10g, Sigma-Aldrich), and flow cytometry for the identification of GFP expression was performed within two weeks of transduction.

### 4.14 Scratch Migration Assay

Cisplatin sensitive and resistant PDX4 cells were seeded in 48-well tissue culture plates, with 1.0x10^6^ and 1.2x10^6^ cells per well, respectively, according to their growth rate and phenotypes to achieve overnight confluence. Once the plates reached confluency, growth media was replaced with FBS-free media, and serum starvation was conducted for 24 hours before wells were scratched with a 10-μL pipet tip. Scratches were imaged at regular time intervals using a Nikon Eclipse Ti microscope. Images were analyzed with ImageJ software.

### 4.15 Colony Formation Assay

PDX4 SE and PDX4 CR cells were seeded at low density in 48-well plates (20 cells/well; n=24 per plating; 25-108MP; Genesee Scientific) for 10 days. On day 5, additional media was added to replenish nutrients. On the final day, surviving colonies were washed with PBS, fixed with methanol mixed with acetic acid (A6283-2.5L; Sigma-Aldrich) solution (3:1 ratio) for 5 to 10 mins, and washed with PBS again before staining. Colonies were stained with 0.1% crystal violet dissolved in methanol and diluted in PBS (1:10) for 20 mins. Crystal violet solution was removed, and residual solution was washed gently with tap water. Colonies were airdried overnight and imaged with an EPSON Perfection V500 Scanner. Colonies were quantified manually and with the ImageJ software automated colony counting feature. Identical parameters were utilized for all the wells.

### 4.16 Statistical Analysis

For all experiments, samples in the same treatment group were harvested from at least three biological replicates and processed individually. All values in the figures and text are the means ± SD. Graphs were generated, and statistical analyses were performed using Prism v 10.1.1. Statistically significant differences were determined by unpaired t-test or two-way ANOVA, unless otherwise noted. P-values less than 0.05 were considered significant.

## ACKNOWLEDGMENTS

We thank Dr. Tuan Zea Tan for assisting with the ovarian cancer molecular subtyping of RNA-seq samples, Pedro Ochoa for helping with the assessment of RNA quality for RNA-seq sample preparation, Dr. Greisha L. Ortiz-Hernández for her technical support with the colony formation assay, Ann Morcos for aiding with the cell cycle flow cytometry assays, and Jacqueline Coats for her technical support and expertise in flow cytometry panel design and optimization. We thank the NCI-funded BigCare Training for Cancer Research workshop hosted by Purdue University for providing training as well as access to their servers, enabling us to analyze the RNA-seq data. We are also grateful to Drs. Carlos A. Casiano and Salvador Soriano for their helpful discussions in topics of mechanisms of cell death. This work was funded by Grant to Promote Collaboration and Translation from Loma Linda University, grant number 2210400 to JJU and YJI.

## ETHICS APPROVAL AND CONSENT

Patients gave their informed consent before participation in the study.

## AUTHOR CONTRIBUTIONS

Conceptualization, T.S. and J.J.U.; Experimental designs, T.S., A.C., Y.J., R.B., A.A., W.C., Y.J.I., and J.J.U.; Data collection, T.S., A.C., Y.J., R.B., A.A., and W.C.; Formal analysis,T.S., A.C., Y.J., and G.Y.; Data interpretation, T.S., A.C., Y.J., and J.J.U.; Project administration,T.S. and J.J.U.; Supervision, J.J.U. and C.W.; Validation, T.S., A.C., Y.J.; Writing, T.S. and J.J.U. All authors revised the manuscript.

## DATA AVAILABILITY

The raw gene expression data and code that support the findings of this study are available from the corresponding author upon reasonable request.

