## Supplemental files for "Modeling platinum resistance in a stem-like patient-derived ovarian cancer sample"

**Supplementary Table 1**. Primer sequences for RT-qPCR.

| Gene | Forward Primer Sequence (5' – 3') | Reverse Primer Sequence (5' – 3') |
|---|---|---|
| ***ACTB*** | TGA AGT GTG ACG TGG ACA TC | GGA GGA GCA ATG ATC TTG AT |
| ***CD44*** | TGG CAC CCG CTA TGT CCA | GTA GCA GGG ATT CTG TCT G |
| ***FN*** | GCC TAA GCA CTG GCA CAC AAC AGT TT | ACT GCC ATT GAT GCA CCA TCC AAC |
| ***LIN28A*** | GAG CAT GCA GAA GCG CAG ATC A | TAT GGC TGA TGC TCT GGC AGA A |
| ***NANOG*** | CAA AGG CAA ACA ACC CAC TT | TCT GCT GGA GGC TGA GGT AT |
| ***POU5F1*** | AAG CGA TCA AGC AGC GAC TAT | GGA AAG GGA CCG AGG AGT ACA |
| ***SNAI1*** | CAC TAT GCC GCG CTC TTT C | GGT CGT AGG GCT GCT GGA A |
| ***SNAI2*** | AAG CAT TTC AAC GCC TCC AAA | AGG ATC TCT GGT TGT GGT ATG AC |
| ***SOX2*** | GCG CCC TGC AGT ACA ACT C | GCT GGC CTC GGA CTT GAC |
| ***TWIST1*** | GGA GTC CGC AGT CTT ACG AG | TCT GGA GGA CCT GGT AGA GG |
| ***ZEB2*** | TTC CTG GGC TAC GAC CAT AC | TGT GCT CCA TCA AGC AAT TC |

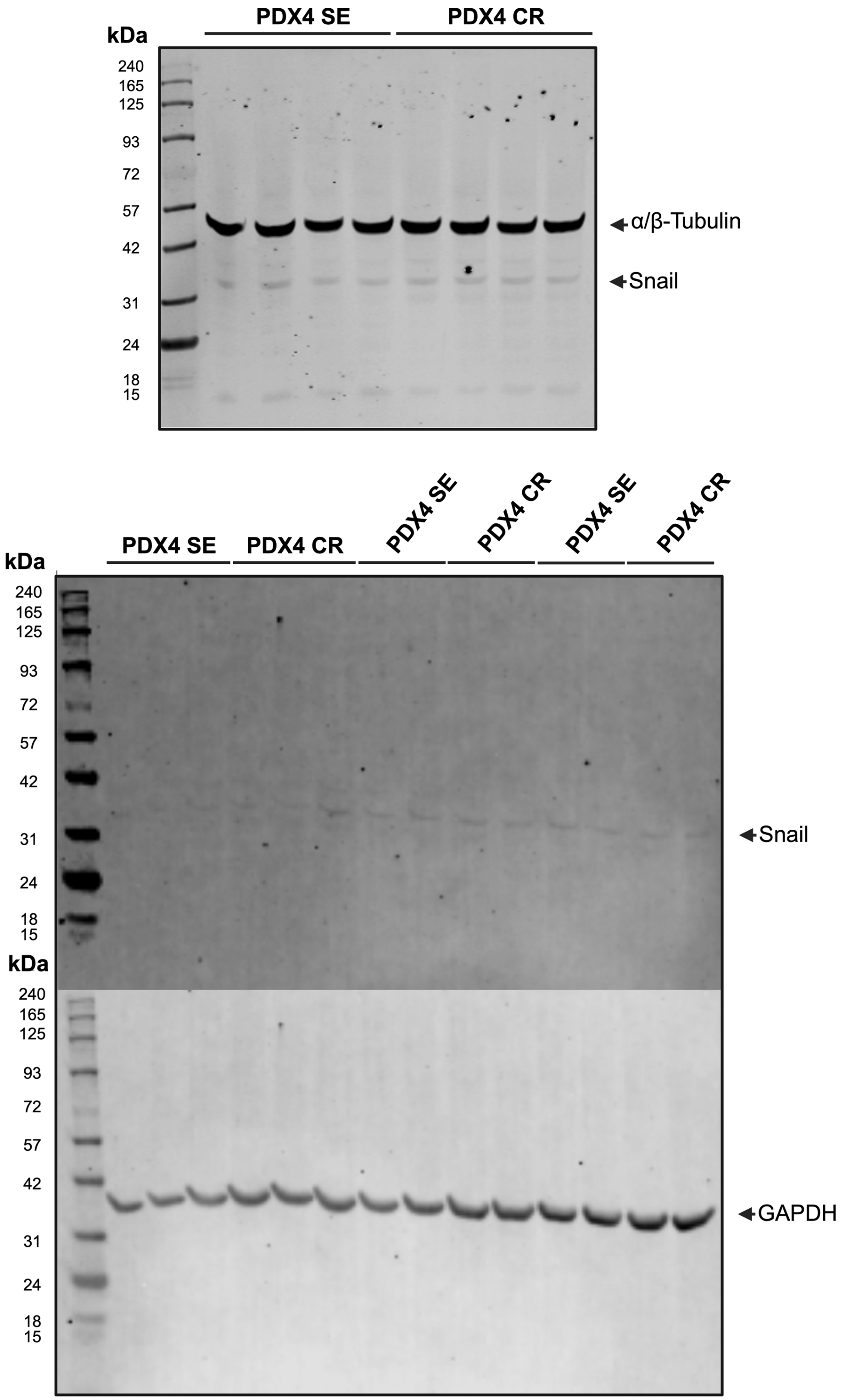


**Supplementary Figure 1. Snail Western blots.** Whole protein was extracted from PDX4 SE and CR. After protein quantification with BCA protein assay, fifty micrograms of protein was added per lane and electrophoresed for 1 hr and 50 min. Protein transfer to PVDF membranes was done using Genscript's eBlot$^{TM}$ L1 Fast Transfer System with the manufacturer's recommended settings. Membranes were then probed with anti-Snail and anti-Tubulin/Gapdh (internal controls), washed, and scanned with Licor Odyssey DLx Imager.

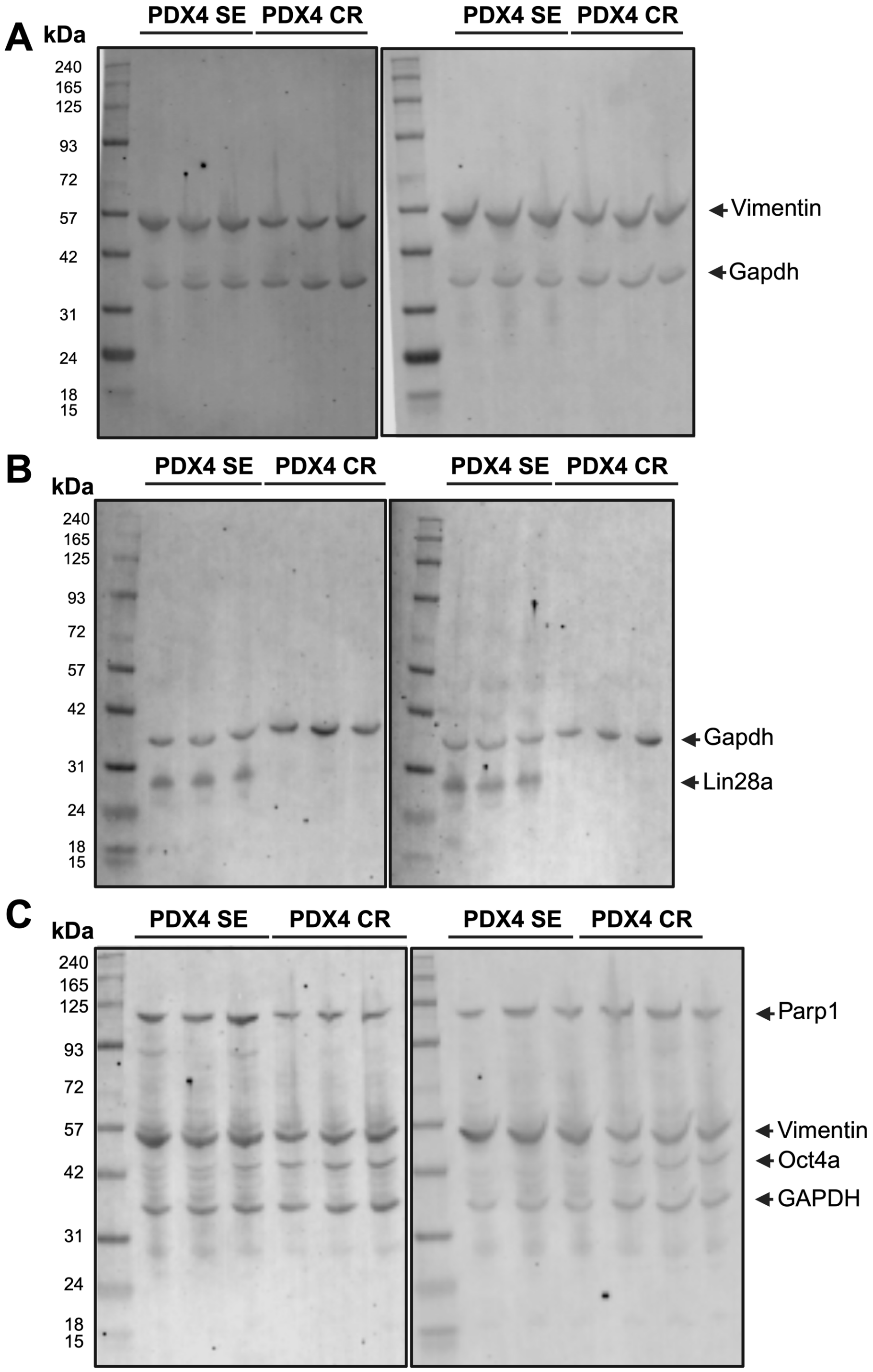


**Supplementary Figure 2. Vimentin, Lin28a, Parp1, and Oct4a western blots.** Whole protein was extracted from PDX4 SE and CR. After protein quantification with BCA protein assay, fifty micrograms of protein was added per lane and electrophoresed for 1 hr and 50 min. Protein transfer to PVDF membranes was done using Genscript's eBlot$^{TM}$ L1 Fast Transfer System with the manufacturer's recommended settings. Membranes were then probed with anti-Vimentin (**A**), anti- Lin28a (**B**), anti-Parp1, anti-Oct4a (**C**), and anti-Gapdh (internal control), washed, and scanned with Licor's Odyssey DLx Imager.

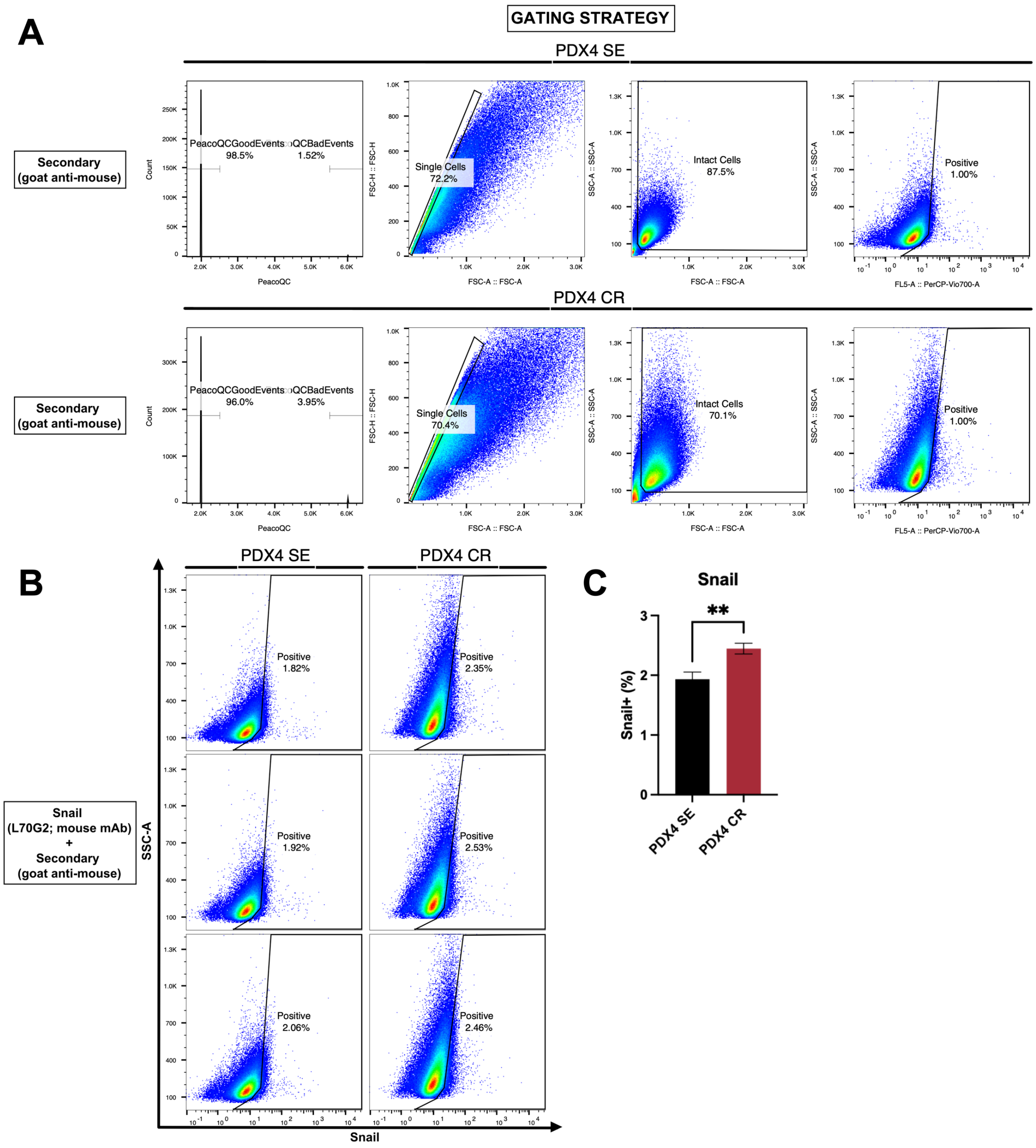


**Supplementary Figure 3. Quantification of Snail expression with flow cytometry. A.** Gating strategy for the identification of positive population relative to populations stained with only secondary antibody. **B.** Dot plots demonstrating proportion of Snail positive cells. **C.** Bar chart depicting PDX4 CR with a significantly higher expression of Snail when compared to PDX4 SE.

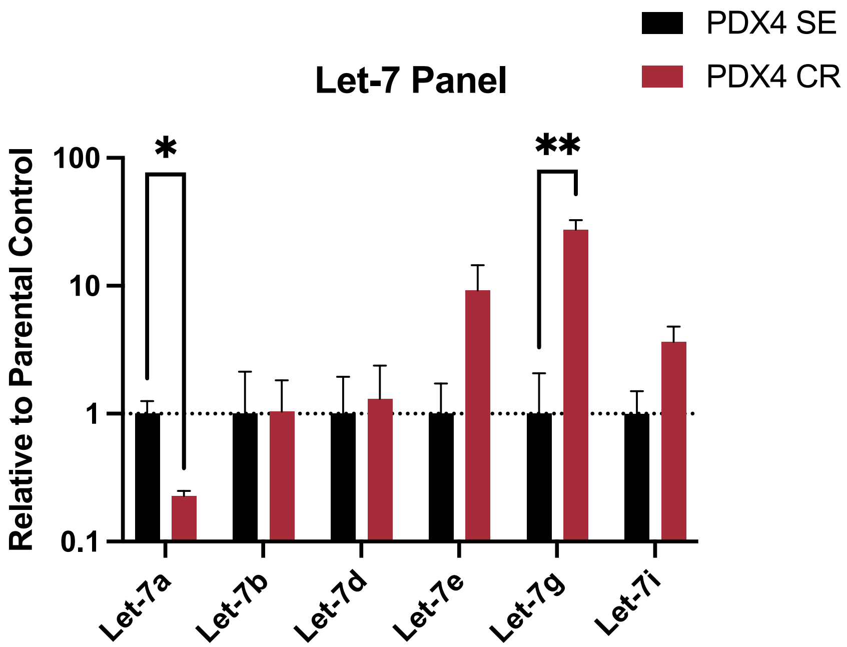


**Supplementary Figure 4. Expression of Let-7 family members in cisplatin sensitive and resistant cells.** RT-qPCR quantification of mature miRNAs.

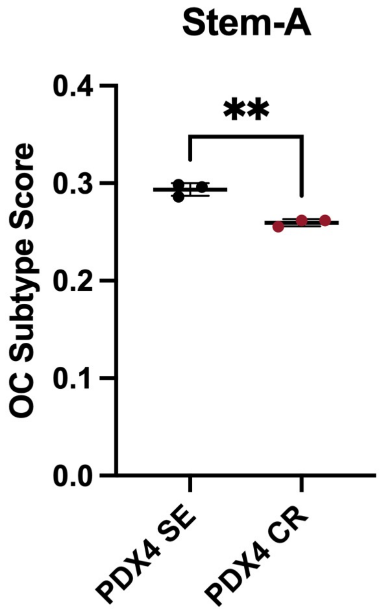


**Supplementary Figure 5. Ovarian cancer molecular subtype classification.** PDX4 SE has a stronger Stem-A signature compared to PDX4 CR.

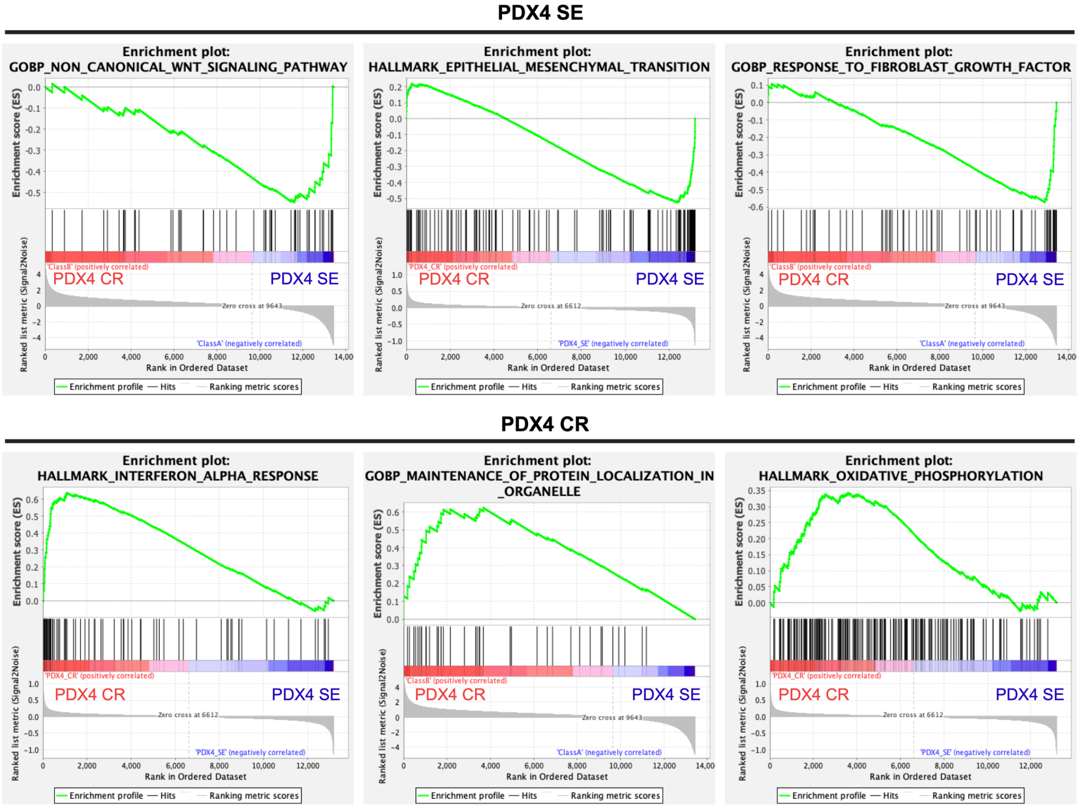


**Supplementary Figure 6. Gene set enrichment analysis for cisplatin sensitive and resistant PDX4.** PDX4 SE had an enrichment in genes associated with non-canonical Wnt signaling, epithelial-mesenchymal transition, and response to fibroblast growth factor. PDX4 CR had an enrichment in genes associated with interferon alpha response, maintenance of protein localization in organelle, and oxidative phosphorylation.

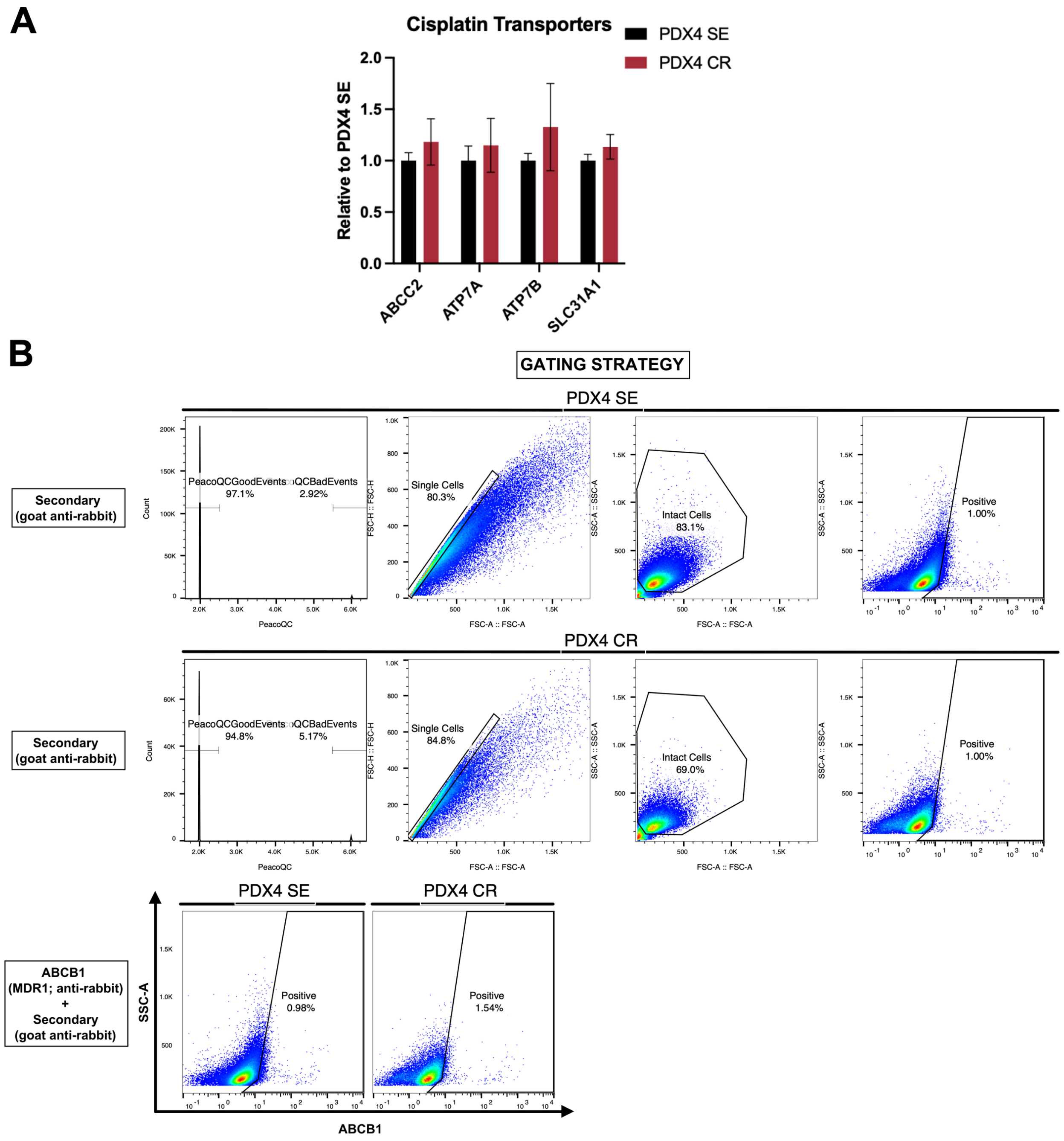


**Supplementary Figure 7. Quantification of cisplatin/copper transporter expression. A.** RT-qPCR quantification of ABCC2, ATP7A, ATP7B, SLC31A1. **B.** Gating strategy and quantification of ABCB1/MDR1 through flow cytometry.
